# deepBlastoid: A Deep Learning-Based High-Throughput Classifier for Human Blastoids Using Brightfield Images with Confidence Assessment

**DOI:** 10.1101/2024.12.05.627041

**Authors:** Zejun Fan, Zhenyu Li, Yiqing Jin, Arun Pandian Chandrasekaran, Ismail M. Shakir, Yingzi Zhang, Aisha Siddique, Mengge Wang, Xuan Zhou, Yeteng Tian, Peter Wonka, Mo Li

## Abstract

Recent advances in human blastoids have opened new avenues for modeling early human development and implantation. Human blastoids can be generated in large numbers, making them suitable for high-throughput screening, which often involves analyzing vast numbers of images. However, automated methods for evaluating and characterizing blastoid morphology are still underdeveloped. We developed a deep-learning model capable of recognizing and classifying blastoid brightfield images into five distinct quality categories. The model processes 53.2 images per second with an average accuracy of 87%, without signs of overfitting or batch effects. By integrating a Confidence Rate (CR) metric, the accuracy was further improved to 97%, with low-CR images flagged for human review. In a comparison with human experts, the model matched their accuracy while significantly outperforming them in throughput. We demonstrate the value of the model in two real-world applications: (1) systematic assessment of the effect of lysophosphatidic acid (LPA) concentration on blastoid formation, and (2) evaluating the impact of dimethyl sulfoxide (DMSO) on blastoids for drug screening. In the applications involving over 10,000 images, the model identified significant effects of LPA and DMSO, which may have been overlooked in manual assessments. The deepBlastoid model is publicly available and researchers can train their own model according to their imaging conditions and blastoid culture protocol. deepBlastoid thus offers a precise, automated approach for blastoid classification, with significant potential for advancing mechanism research, drug screening, and clinical in vitro fertilization (IVF) applications.

## Introduction

Mammalian development begins with a fertilized egg that has the potential to form all embryonic^1, 2, 3^ and extra-embryonic lineages^4, 5, 6^. As development progresses, cells within the embryo gradually lose their potency and begin to differentiate into specialized cell types^7^. In the human embryo, the earliest cell fate decisions are made by the time of blastocyst formation^8^, just prior to implantation.

The blastocyst, which emerges approximately three and a half days post-fertilization (E3.5) in mice and around five days in humans, is defined by a fluid-filled cavity and consists of three distinct cell populations^9^: the outer extra-embryonic trophectoderm (TE), and the inner epiblast (EPI) and primitive endoderm (PrE), which are positioned adjacent to the cavity. Due to ethical considerations of experimenting with human embryos, alternative models have been developed to recapitulate many aspects of the human peri-implantation development^10, 11^. Recent studies^12, 13, 14, 15, 16, 17, 18^ have demonstrated that stem cells can self-organize in vitro to generate blastocyst-like structures(Fig. 1a), termed blastoids. By closely mimicking early human embryonic stages, blastoids offer an ethical and cost-effective alternative for understanding the mechanisms underlying embryogenesis, investigating pregnancy-related diseases such as abnormal development and pregnancy failure, and testing drug safety and efficacy in a controlled environment^19, 20^.

**Fig. 1.**
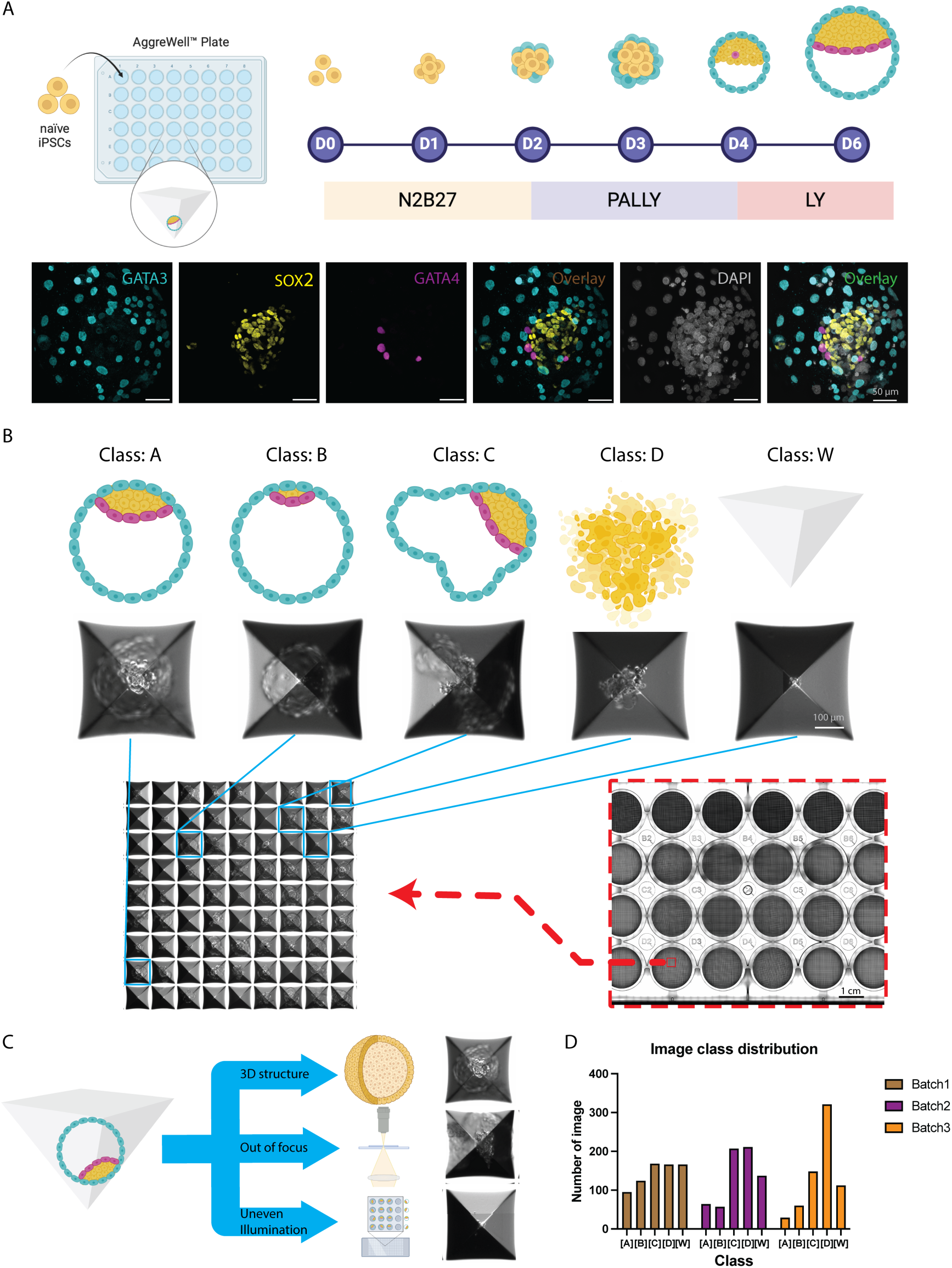
Generation of a blastoids image dataset for our classification model. (**A**), Protocol of blastoid cluture and Immuno-fluorescence microscopy images of blastoids. The blastoids were fluorescently labelled with GATA3 antibody (magenta), SOX2 antibody (yellow), GATA4 antibody (cyan) and DAPI (gray). Scale bar, 50 μm. (**B**), Five typical classes of blastoids generated in a high-throughput microwell. Top: schematic of different classes of blastoids. Middle: brightfield images of blastoids in microwell. Scale bar, 100 μm. Bottom: generation of blastoids in AggreWell plate. Each plate has 24 wells. Each well has ∼1200 individual microwells for blastoids formation. Scale bar, 1 cm. (**C**), Schematic of three common interferences hindering traditional image analysis in blastoids classification. (**D**), Distribution histogram of blastoids images in three individual batches, evaluating the uniformity of dataset.

Over the past year, various methods^12, 14, 16, 17, 21^ for generating human blastocyst models have been proposed, with most being implemented in the Aggrewell format^16, 22^. However, the automatic characterization of blastoids remains largely underdeveloped. Critical parameters, such as cavitation efficiency, are still manually evaluated by researchers and large amounts of experimental data remain underexplored, leading to missed opportunities for deeper insights^23, 24^. The absence of automated, comprehensive characterization tools often results in valuable information being overlooked^25^. Traditional image segmentation tools are insufficient for blastoid images analysis due to the compression of their 3D-structure into a single 2D-image, compounded by issues like variations in focal plane and uneven illumination (Fig. 1c). These challenges are common and unavoidable in brightfield images of blastoids in Aggrewell, increasing the burden of manual evaluation and limiting their application in drug screening and mechanistic research.

To address these challenges, we developed a deep-learning classifier based on a fine-tuned ResNet-18^26, 27, 28^ model for high-throughput identification and classification of blastoids. This model, using only brightfield images, is capable of automatically classifying blastoids into five distinct quality categories. The trained classifier processes an average of 53.2 images per second with an accuracy of 87%, without signs of overfitting or batch effects. To further enhance accuracy, we introduced a Confidence Rate (CR) mechanism, which flags low-confidence classifications for human expert review, thereby increasing the overall accuracy to 97%, albeit with a slight reduction in throughput. We demonstrate two real-world applications of this deep-learning model using wet lab experiments: 1) analyzing the effect of lysophosphatidic acid (LPA) dosage gradients on blastoid formation across different classes, and 2) assessing the impact of DMSO on blastoids for drug screening. Our findings indicate that the minimum effective concentration (MEC) of LPA is 0.5 µM, which notably promotes the formation of Class B blastoids. Meanwhile, DMSO, a commonly used pharmaceutical solvent^29, 30^, showed no effect on blastoid formation when used at a 0.1% concentration compared to the control group. Additionally, the model facilitated the assessment of cell seeding density as a quality control measure, thereby strengthening the reliability of our conclusions.

In summary, this deep-learning model provides a detailed and automated classification method for evaluating blastoids, uncovering new insights into developmental mechanisms^31,32^. For drug screening^33, 34, 35^, the high-throughput classifier offers an efficient, accurate, and easy-to-use solution, enabling large-scale screening of blastoids. Moreover, with transfer learning^36, 37^, this model could potentially be applied in clinical settings, such as in vitro fertilization (IVF), to improve blastocyst evaluation^38, 39, 40^ and reduce pregnancy failure rates.

## Results

### Establishment of a blastoid dataset

As illustrated in Fig. 1a, over six days following the seeding of naïve iPSCs in the AggreWell plate, the cells proliferated, differentiated, cavitated, and eventually formed blastoids. Brightfield images of each blastoid in every microwell were captured under a microscope to create the blastoid image dataset. The formed blastoids exhibit a characteristic cavity structure and three cell types—SOX2^+^ EPI, GATA3^+^ PrE, and GATA4^+^ TE—which will develop into the embryo proper, yolk sac, and placenta in vivo, respectively (Fig. 1a). Each AggreWell plate contains 24 wells, with each well housing 1,200 microwells, resulting in nearly 30,000 blastoids per experiment awaiting evaluation (Fig. 1b). These blastoids are classified into five distinct classes (Fig. 1b). Class A represents a blastoid with a well-formed cavity and an well-sized ICM embedded in a complete circular TE layer. Class B represents a blastoid with a cavity and a small or absent ICM, but with a complete circular TE layer. Class C represents a blastoid with an adequate ICM but with an irregular or broken TE layer. Class D represents a failed blastoid that does not form a cavity, appearing as cellular debris. Class W represents an empty microwell with no cells seeded.

Based on class features, human experts carefully and rigorously classified and labeled each image into one of the five defined classes. The class distribution of three independently produced batches of blastoids is consistent, showing similar proportions (Fig.1d). Nevertheless, we observed that the distribution is imbalanced, with an insufficient ratio of Class A blastoids. To deal with the imbalance, we choose to adapt the weights in the loss function in training^41, 42, 43^.

### Architecture of deep learning model

The overall data flow is outlined in the Fig. 2b. Considering the balance between computational efficiency and effectiveness, we choose ResNet18 as our default classification model in this work. As shown in Fig. 2c, given an input image, the model extracts feature maps through successive convolutional layers with shortcut connections (*i.e.*, residuals) and finally projects the last feature to the classification probability via an average pooling, a fully connected layer and a softmax function(Supplementary Table1). These residual connections address the vanishing gradient problem, enabling the training of such deeper networks by allowing gradients to flow more effectively through the network during backpropagation. The output probability indicates the classification confidence that can be further utilized in the human review process.

**Fig. 2.**
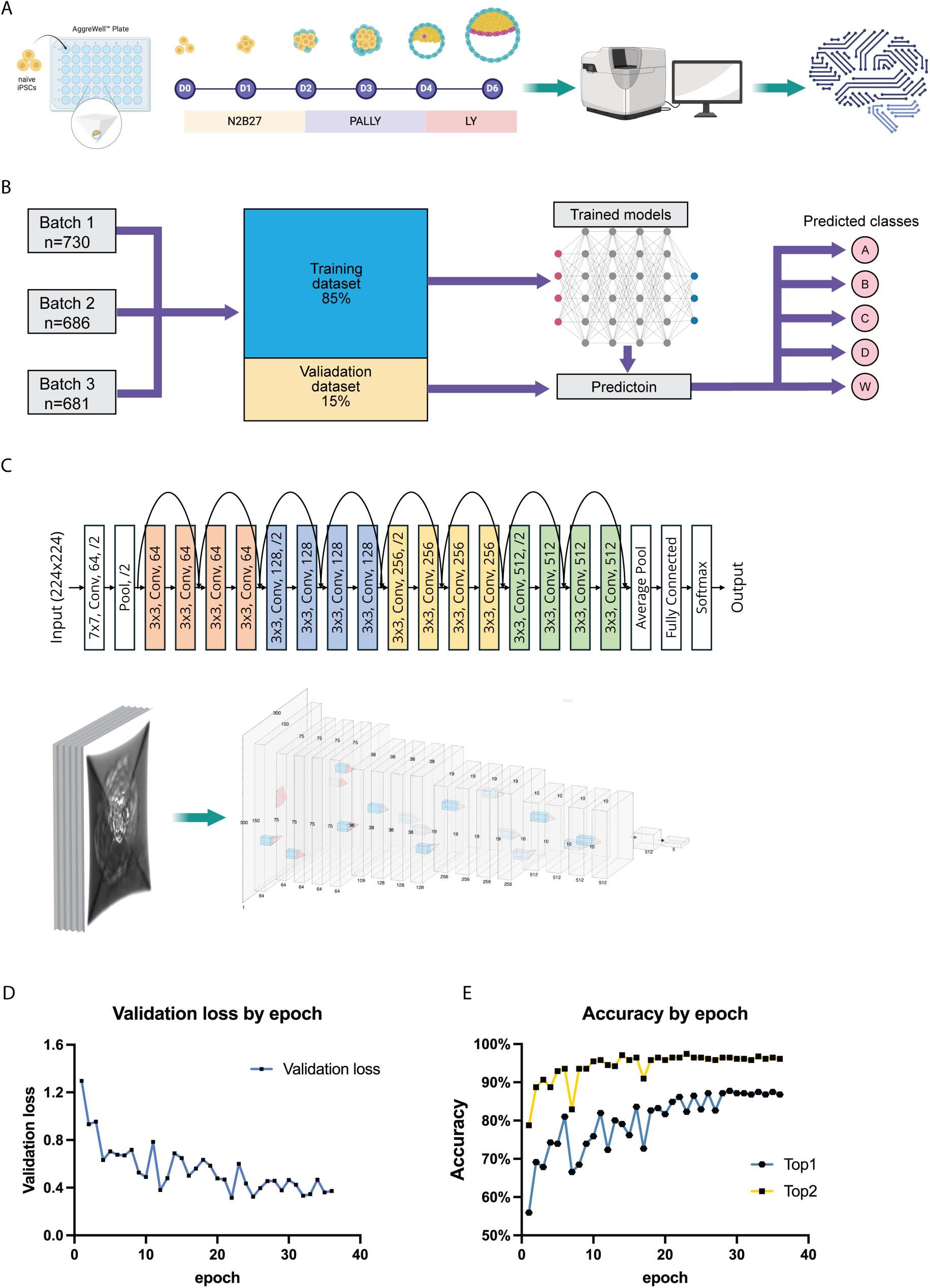
Training of blastoids classification model. (**A**), Schematic representation of the assay workflow used to generate the original blastoid images. The cells are seeded in the AggreWell Plate, cultured in different medium and harvested at Day6. Images are acquired by CellDiscoverer7 microscopy, followed by AI model training. (**B**), Workflow of the model. All images form three batches are evenly divided into training dataset (85%) and validation dataset (15%). Using the training dataset, the model is cross-trained and followed by validation. (**C**), Architecture of deep learning ResNet18. The architecture is composed of seventeen convolutional layers, one max pooling layer, one average pooling layer and one fully connected layers. Representative loss curve (**D**) and Top1/Top2 accuracy curves (**E**) for the validation dataset versus epoch in training. The loss curve and accuracy curves are stable after epoch 36.

### Data processing and model training

We used the three batches of blastoid data consisting of 2107 images, and randomly divide them into a training set with 1796 data pairs (85%) and a validation set with 311 data pairs (15%), respectively. Then, we further adopt 728 labeled data pairs and 14726 unlabeled images as the test set for model evaluation.

To diversify the training data and further improve the model performance and generalization, we adopt various data augmentation strategies during the training stage, including horizontal flip, vertical flip^44^, color jitter^45^, and random erasing^46^. The probability for adopting color jitter and random erasing is set to 40% and 20%, respectively.

We employ the cross-entropy loss and AdamW optimizer to train our model for 36 epochs with a learning rate of 3 ∗ 10^-4^ and a weight decay of 1 ∗ 10^-3^. We clip the gradients to stabilize the training and scale the learning rate of the last fully connected layer with 10 to achieve faster convergence. As shown in Fig. 2e, the validation loss decreases with some volatility during training and eventually plateaus after 24 epochs, indicating the point at which the model is well-trained. Similarly, both Top-1 and Top-2 accuracy on the validation set follows a similar trend reaching the plateau after 24 epochs. We selected the model trained with 36 epochs to ensure convergent.

### Performance of the deep learning classifier

The confusion matrix of the validation dataset, which displays the number of correct and incorrect predictions for each class to show how well the model has identified each category, illustrates the performance of the deep learning classifier (Fig. 3a). Most of the data points are concentrated along the diagonal, which is shaded in dark blue, indicating strong prediction and classification accuracy. However, some light blue areas appear near the diagonal, signifying misclassifications, particularly between Class A and Class C. These two classes share similar biomorphological features, which likely results in the misclassification. Overall, the classifier achieved a Top-1 accuracy of approximately 87% and a Top-2 accuracy of 97% (Fig. 3b). Furthermore, the model demonstrated nearly equal accuracy across the three experimental batches (Fig. 3b), indicating good generalizability and no signs of overfitting or batch-effects during the training process.

**Fig. 3.**
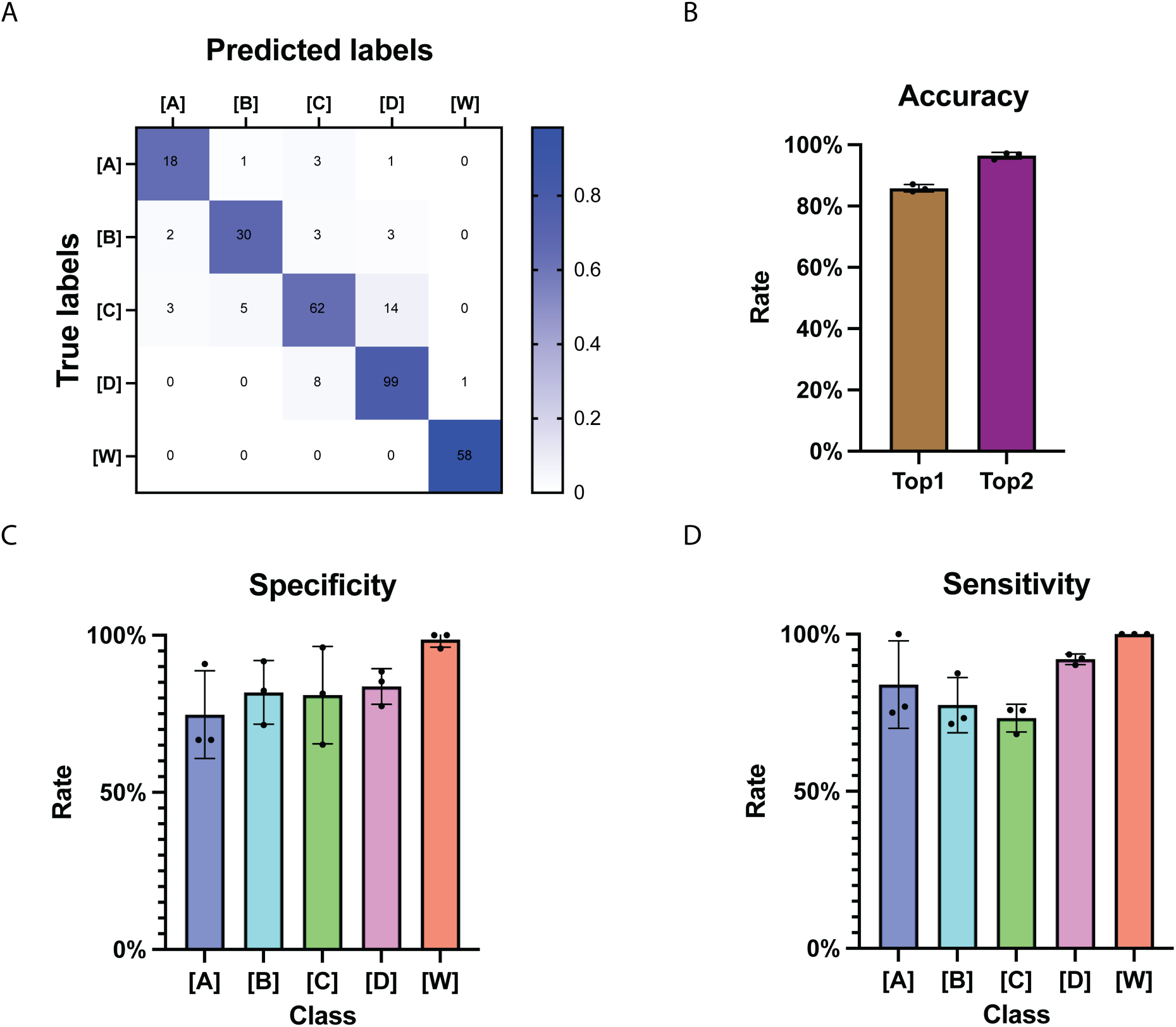
Performance of the blastoids classification model. (**A**), Confusion matrix on the test set. The color in cells illustrates distribution levels. The number in cells shows each number of images. (**B**), Top1/Top2 accuracy of model. Specificity (**C**) and Sensitivity (**D**) of model for five classes. Each point is from an individual batch dataset.

More detailed metrics, such as specificity and sensitivity, are provided in Fig. 3c and 3d to further assess the model’s performance. Specificity values remain consistent across the three batches, with no significant differences among classes and values close to the average accuracy, indicating that the model is well-trained. The AI model performs best on Class W, as shown in the confusion matrix in Fig. 3a. Sensitivity also demonstrates stable performance across the three batches, although Class C has a lower sensitivity than the others. Together, these metrics provide a comprehensive evaluation of the model’s performance.

### Confidence rate enhanced classifier

While the classifier achieved adequate accuracy, we sought to further reduce misclassification and improve overall performance. To achieve this, a confidence rate (CR) was introduced to filter out images that are likely to be misclassified, which would then be reviewed by human experts (Fig.4a). For each input image, the model outputs probabilities for all potential classes, which are then post-processed. The CR is calculated based on the integration of these probability features.

As shown in Fig. 4a, there are two stages in the data flow where human involvement is critical: defining the labels and reviewing images with low CR. Fig. 4b illustrates that a trained classifier was used to generate the initial labels, while CR features were provided on the fly, assigning a probability to each class for further processing.

**Fig. 4.**
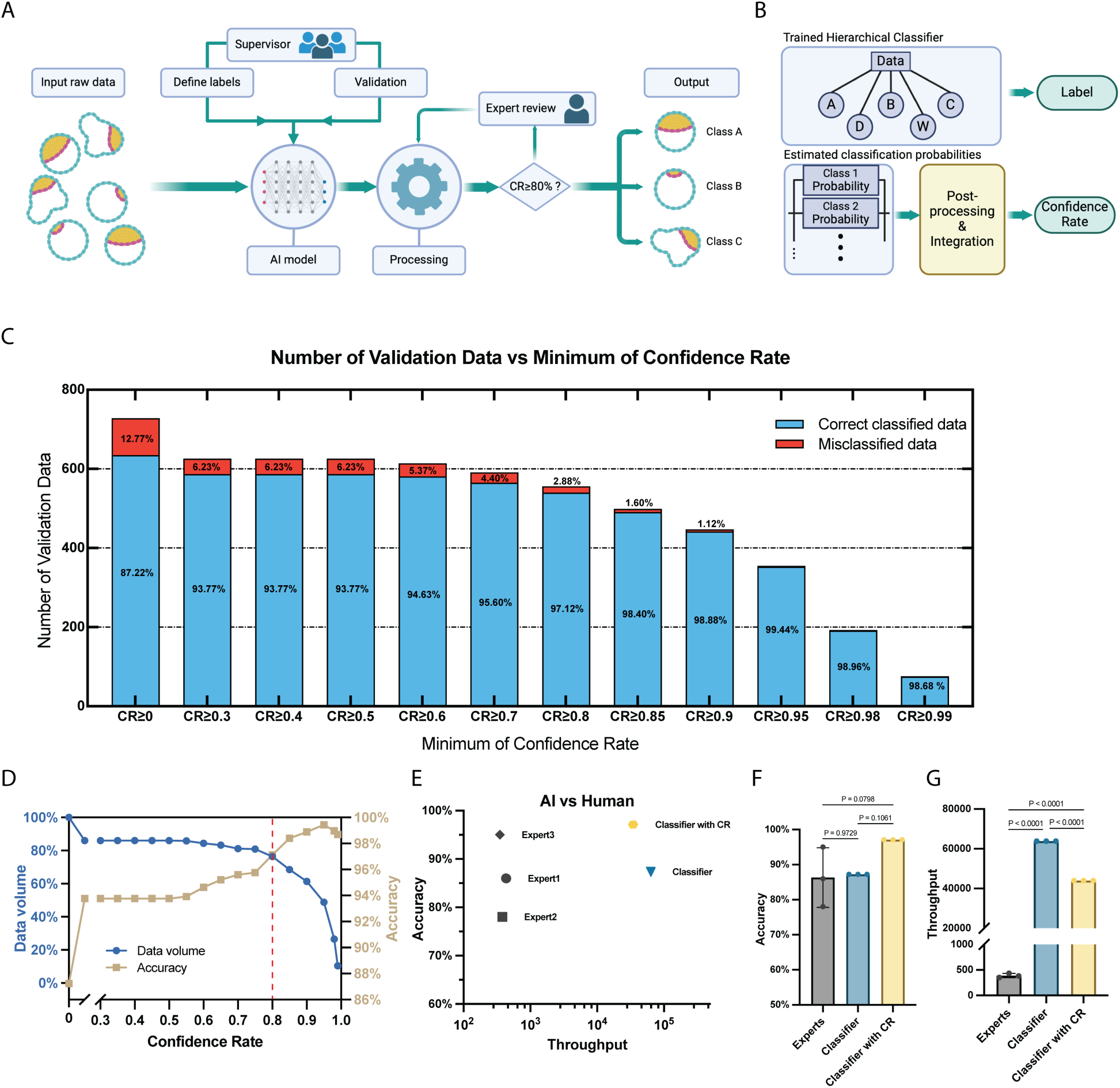
Classifier of blastoids images with confidence assessment. (**A**), Data flow of classifier with confidence rate and human expert review. (**B**), Schematic of proposed classifier with confidence rate. The trained hierarchical classifier selectively utilizes features for labeling purposes. A misclassification probability is assigned to each feature using the estimated Probability Density Functions (PDFs). These probabilities are aggregated to a confidence rate for the concluded label. (**C**), The performance of the proposed classifier with confidence rate on an independent dataset. (**D**), Relationship between minimum CR and Data Volume & Accuracy. Beige curve is for accuracy. Blue curve is for data volume. Red dot line is for the intersection where CR=0.8. (**E**), Comparison of performance between Deep learning model and human experts in throughput and accuracy. The x-axis is for the number of images processed by AI or human in 20 mins. In the Classifier with CR, 0.8 is chosen as an acceptable confidence rate. (**F**), Quantitative analysis of accuracy performance of human experts and Deep learning model. (**G**), Quantitative analysis of throughput of human experts and Deep learning model.

We then define a CR threshold to select unreliable samples for reviewing by a human expert. As CR increases, the amount of misclassified data decreases, and accuracy improves from 87% to 99% (Fig.4c). However, this comes with a significant reduction in data volume, placing a heavier burden on human reviewers and diminishing the classifier’s practicality. Therefore, a balance between data volume and accuracy must be achieved. As shown by the yellow accuracy curve and blue data volume curve in Fig. 4d, a CR value of 0.8 was selected as the optimal minimum CR. This value represents a point before the data volume drastically declines while accuracy significantly increases from 87% to 97%, as indicated by the red line in Fig. 4d.

### Comparison with human experts

To compare the performance in terms of throughput and accuracy, we conducted a competition between the AI classifier and human experts on the test set, which showed that the AI classifier demonstrated extremely high processing throughput and comparable accuracy to that of the three human experts (Fig. 4e). Additionally, the classifier, when combined with the confidence rate (CR) feature, leveraged the strengths of both human experts and AI, maintaining high accuracy while achieving acceptable throughput levels.

More specifically, the AI classifier shows no significant difference in accuracy compared to the human experts (Fig. 4f), highlighting the classifier’s reliability. Meanwhile, the AI classifier’s throughput is significantly higher—by several orders of magnitude—than that of human experts (Fig. 4e), emphasizing its efficiency and scalability.

### Field application I: LPA dosage optimization for blastoid formation

During blastoid formation, various inhibitors and growth factors are used to induce differentiation of different cell lineages that resemble the TE, EPI and PrE in blastocysts. The concentrations of these factors require careful titration in protocol development and optimization when using different cell lines. However, systematic optimization is time-consuming and labor-intensive, meaning it is often shortened in real-world applications, resulting in suboptimal conditions. One of such factors, lysophosphatidic acid (LPA), a key compound in the PALLY medium^22^, plays a crucial role in inducing blastoid formation^47, 48^. LPA acts through its receptors to regulate various cellular processes, including the inhibition of the Hippo pathway during TE development^49, 50^(Fig. 5a). To investigate the effect of different LPA dosages on blastoid formation efficiently, we performed blastoid generation experiments and utilized the model to analyze the results, significantly reducing the labor and time required for manual analysis. A gradient of LPA doses, including 0, 0.5, 1, 2.5, and 5 µM, was tested, and over 10,000 images from three replicate batches were processed by the classifier (Fig. 5a). Consistent with previous studies^51^, cavitation efficiency increased with higher LPA dosages, with the minimum effective concentration (MEC) being 0.5 µM. Beyond 0.5 µM, no significant differences were observed in cavitation efficiency (Fig. 5b).

**Fig. 5.**
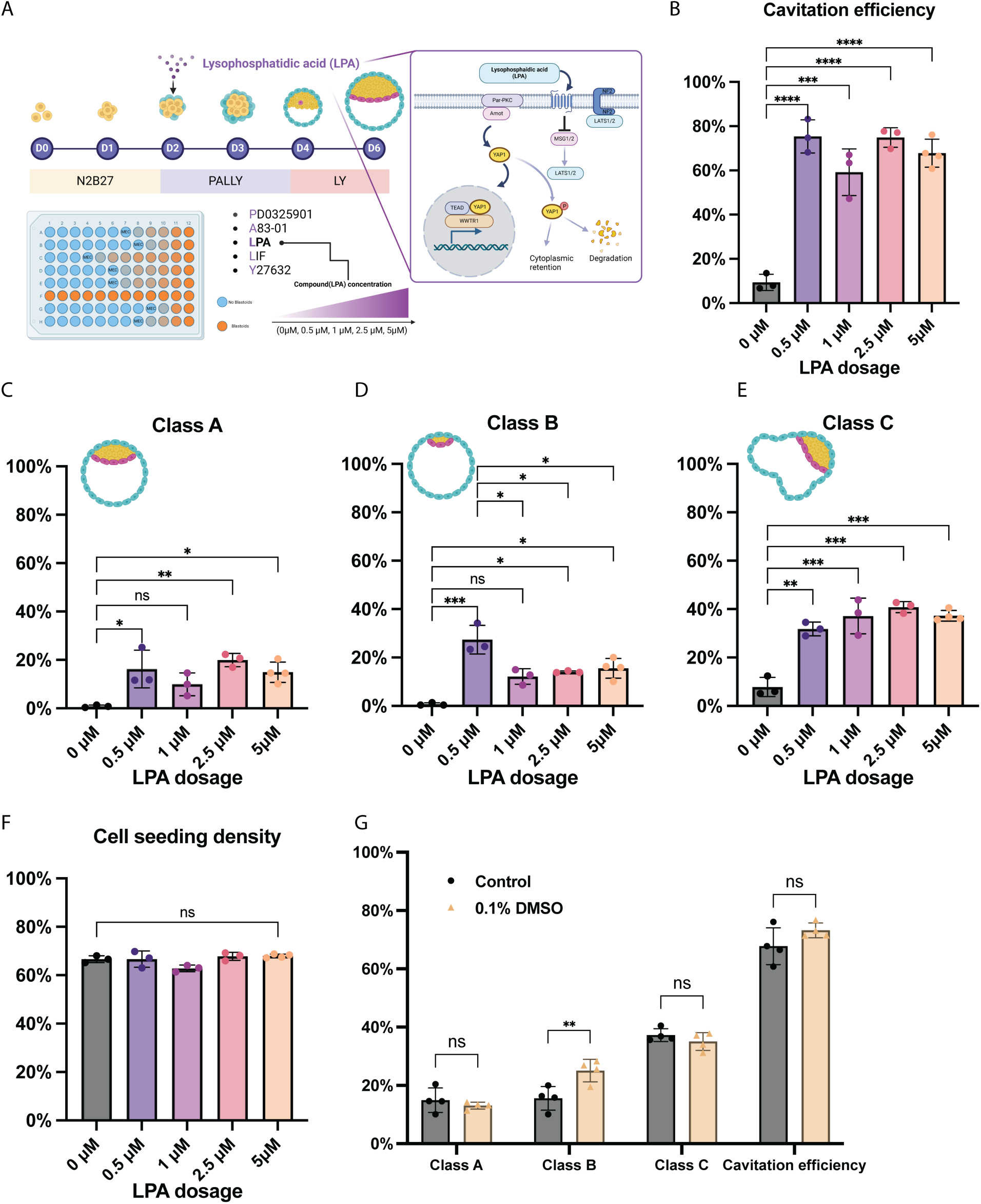
Application of classifier model for blastoids. (**A**), Schematic of dose-effect experimental design of LPA on blastoids formation, including 0 μM, 0.5 μM, 1 μM, 2.5 μM and 5 μM. All culture conditions are normal, except LPA concentration. Purple box indicates a diagram about LPA and its role in blastoid formation by inhibiting the Hippo pathway. LPA has a significant dose-effect on blastoids formation in the aspect of cavitation efficiency (**B**), class-A efficiency (**C**), class-B efficiency (**D**), class-C efficiency (**E**). (**F**), Cell seeding density in the gradient dosage experiment. Cell seeing density is the proportion of seeded wells, the sum of class A, B, C, and D. (**G**), Effects of 0.1% DMSO on blastoids formation in different classes. Each point is for an independently replicated experiment. One-way ANOVA and Tukey’s multiple comparisons test. *P < 0.05, **P < 0.01, ***P < 0.001, ****P < 0.0001.

Additionally, similar trends were observed in Class A, B, and C blastoids, as shown in Fig. 5c, 5d, and 5e, respectively. Notably, Fig. 5d revealed a significant two-fold increase in Class B at the 0.5 µM dosage compared with 1uM, 2.5uM and 5uM, a phenomenon not observed at higher dosages. This suggests that 0.5 µM may represent an optimal concentration for blastoid formation. Without the AI classifier’s automated classification, this observation could have been easily overlooked. Further biological experiments are necessary to investigate the underlying mechanisms behind this finding. Moreover, as illustrated in Fig. 5f, no differences in cell seeding density were found between batches across the LPA gradient, indicating that the AI classifier provided strong quality control during the classification process, ensuring the experiment’s reliability.

### Field application II: DMSO eDect on blastoid formation

The AI classifier can also be leveraged to investigate the effect of compounds on blastoid formation. DMSO, commonly used as a solvent in drug screening^29^, is present in low concentrations during blastoid drug screening experiments. However, its specific impact on blastoid formation is only starting to be studied^21^. We hypothesize that by utilizing the AI classifier, this effect can be easily explored. As shown in Fig. 5g, there is no significant difference between the control group and the 0.1% DMSO group in terms of Class A, Class C, and overall cavitation efficiency. Interestingly, the efficiency of Class B blastoids is higher in the 0.1% DMSO group, suggesting that DMSO may promote trophectoderm (TE) cell formation. Overall, low-concentration DMSO does not appear to affect blastoid cavitation formation, consistent with a previous study^21^.

## Discussion

In conclusion, to efficiently and comprehensively evaluate blastoids, we developed a deep-learning classifier based on a ResNet-18 model for high-throughput identification and classification. Using only brightfield images, the model automatically classifies blastoids into five distinct quality-level classes. After training, this classifier processes 53.2 blastoid images per second with 87% accuracy, showing no signs of overfitting or batch effects. To further minimize misclassification, we incorporated a Confidence Rate (CR) mechanism, where images with low CR scores are flagged for human review, thus ensuring accurate decisions. This combination of AI and human expert raised accuracy from 87% to 97%, though it reduced processing throughput. Two key field applications were demonstrated using this deep-learning model: (1) the effect of a gradient dosage of LPA on blastoid formation, and (2) the effect of DMSO on blastoid formation for drug screening. We found that 0.5 µM significantly promotes Class B blastoid formation. Additionally, 0.1% DMSO had no significant effect on blastoid formation compared to the control group. Importantly, the AI model also tracked cell seeding (Fig. 5f), providing a robust quality control that strengthens the reliability of these experiments. Therefore, this deep-learning model demonstrates high accuracy and throughput, offering significant potential for mechanism research, drug screening, and clinical in vitro fertilization (IVF).

The deepBlastoid model, which is publicly available and adaptable for use beyond our specific system, is designed to be easy-to-use and applicable across diverse research and clinical settings. Researchers can train their own models based on this framework, tailoring it to their specific imaging conditions and blastoid culture protocols, which holds significant value for a wide range of applications. Current studies primarily focus on cavitation efficiency, leaving other valuable information unexamined^12, 21^. For mechanism research, the deep-learning model offers a solution by automatically conducting comprehensive evaluations and detailed classifications of blastoids, potentially revealing new phenomena. In drug screening, this high-throughput classifier is both labor-and time-efficient, facilitating large-scale assessments of blastoids^23^. It offers a promising approach for evaluating drug effects on embryogenesis and pregnancy in pharmaceutical contexts^52^. Clinically, the well-trained model could be adapted to evaluate blastocysts in IVF^53^, thereby reducing the risk of pregnancy failure. Transfer learning from blastoids to blastocysts would also minimize the use of valuable embryos, making the process more ethical and efficient.

However, there are limitations to the project. The dataset used for training, while effective, is relatively small and lacks diversity. Although the model achieves 87% accuracy with thousands of images, this accuracy could be improved with a larger and more varied dataset. The imbalance in the training data, particularly the underrepresentation of Class A, has led to a lower sensitivity in Class A classification. Increasing the number of Class A images would likely enhance the model’s performance. Additionally, to improve the model’s applicability and generalizability, future efforts should involve training with datasets from various imaging conditions^54^, cell lines, and blastoid culture protocols. While the current classification into five classes is based on morphology, a detailed classification into more classes could provide a more comprehensive understanding of blastoid development.

Furthermore, this evaluation is somewhat one-dimensional, focusing solely on morphology. Other important aspects, such as developmental sequences and cell types, should also be included in future evaluation or even prediction by the AI model^55^. For instance, it may become possible to predict cell types and their distribution from brightfield images alone, using related data such as immunofluorescence images for training^56^. If this can be achieved, the deep-learning model would become a powerful non-invasive tool for blastoid evaluation. Moreover, the sequence of blastoid formation contains valuable information that could be captured using time-lapse data and analyzed by the deep-learning model to predict blastoid development. This would be especially valuable for drug screening and mechanistic studies.

In clinical practice, some couples experience pregnancy failure in IVF even when embryos are carefully selected by doctors. The AI model may be able to detect subtle, overlooked information that could aid in evaluating embryos for implantation^38^. Moreover, transfer learning from blastoids to blastocysts may improve the accuracy of embryo selection, reduce the need for valuable embryos, and address ethical concerns. Only fine-tuning for the blastocyst dataset would be required, as the model would already be well-trained on blastoid, offering an efficient pathway for application in clinical settings.

## Materials and methods

### Study design

This study aimed to develop a deep learning model for accurate and high-throughput classification of blastoids based on brightfield images. The design, training, and enhancement of the classifier were key components of the study. The deep learning model was constructed using the ResNet-18 architecture. A dataset of over 2,000 labeled brightfield images of blastoids, generated through established biological protocols, was used to train the model. The performance of the model was evaluated with key metrics including accuracy, specificity, sensitivity, and loss function. After introducing a minimum Confidence Rate (CR) threshold, the model’s accuracy improved significantly, though throughput decreased slightly. A competition between human experts and the AI model further demonstrated the classifier’s superior performance in blastoid classification.

Additionally, the study explored the effect of a gradient dosage of Lysophosphatidic Acid (LPA) on blastoid formation using the deep learning model. In this field application, blastoids were exposed to gradient LPA concentrations, and the model comprehensively evaluated their formation, with cell seeding density serving as a robust quality control measure. In a separate investigation, the model was used to assess the impact of DMSO on blastoid formation by processing image data and evaluating the blastoids. Statistical significance in these analyses was verified using one-way ANOVA.

### Human induced pluripotent stem cells (hIPSCs) and initial culture techniques

This study was reviewed and approved by the KAUST Institutional Biosafety and Bioethics Committee (IBEC). The human pluripotent stem cells (PSCs) used in this study include chemically reset (cR) cR-SC-9N hiPSCs, while primed H1 hPSCs were obtained from the WiCell repository. The primed PSCs underwent epigenetic resetting following an established protocol. Briefly, primed state PSCs were transitioned onto inactivated mouse embryonic fibroblasts (iMEF) and treated with PD0325901 (1 μM), leukemia inhibitory factor (LIF, 10 ng/mL), and valproic acid (VPA, 1 mM) for 3 days. After this initial phase, the cells were transferred to a medium containing 1 μM PD0325901, 2 μM XAV-939, 2 μM GSK-3 inhibitor G.6983, and 10 ng/mL LIF (PXGL). By day 10, naïve dome-shaped colonies began to emerge and were subsequently purified through fluorescence-activated cell sorting (FACS) for SUSD2 or by gelatinization over several passages. The cells were cultured at 37°C under conditions of 5% CO2 and 5% O2.

### Generation of blastoid

To prepare PXGL-naïve pluripotent stem cells (nPSCs) for blastoid differentiation, nPSCs were cultured on inactivated mouse embryonic fibroblasts (iMEF) in PXGL medium for 3–4 days. The cells were then dissociated using Tryple for 5 minutes at 37°C with gentle pipetting until a single-cell suspension was achieved. Tryple was inactivated using 0.2% BSA in N2B27 medium, followed by centrifugation to pellet the cells. To remove any remaining iMEF, the cells were seeded onto a 0.1% gelatin-coated surface for 60 minutes. After this, the cells were passed through a 40 μm strainer to eliminate clumps, centrifuged again, and resuspended in the appropriate medium.

For blastoid formation, we followed an established protocol. Cells were resuspended in N2B27 medium supplemented with a ROCK inhibitor and seeded at a density of approximately 75 cells per microwell in 400 μm AggreWell plates (Stem Cell Technology, cat# 34425). Once the cell structures appeared solid, the medium was changed to N2B27 supplemented with PALLY (1 μM PD03, 1 μM A83, 500 nM LPA, 10 ng/mL LIF, and 10 μM ROCK inhibitor) and maintained for two days or until cavitation was observed. Subsequently, the medium was switched to N2B27 supplemented with LPA (500 nM) and ROCK inhibitor (10 μM) (LY) for an additional two days to allow the structures to mature. The following morphometric criteria were used to define blastoids: blastoids were described as single-layer, cavitated structures with a diameter between 150–250 μm and the presence of a singular inner cell mass.

### Brightfield imaging

Blastoid images were acquired using a Zeiss Cell Discovery 7 microscope (CD7) under brightfield imaging conditions. Given the size of the AggreWell plate, the magnification was set to 10X without additional zoom. The imaging mode was configured as oblique, with an exposure time of 20 ms, using the default light intensity and gain settings. Prior to imaging, the focus was adjusted to capture the largest cross-sectional area of the majority of blastoids. While defocusing is challenging to avoid when capturing large-field tile images, the deep learning model used in this study is robust enough to handle defocused images. To minimize sample shaking, the camera’s velocity and acceleration were both set to 10%.

### Image cropping

After imaging, the AggreWell plate images, which contain thousands of blastoids, were cropped into smaller images corresponding to the size of individual microwells (as shown in Fig 1b) to facilitate subsequent model training. The cropping was performed using the Cropping tool in Photoshop 2023, allowing for precise and efficient segmentation of the large images. No adjustments were made to contrast, brightness, or rotation. The final processed images had a resolution of approximately 300 x 300 pixels, in 16-bit grayscale, and were saved in PNG format.

### Labeling of the dataset

The images were carefully evaluated and labeled by biological experts experienced in embryonic development and blastoid culture. A total of 2,112 images from three experimental batches were classified into five typical categories based on developmental features: Class A, B, C, D, and W, as shown in Fig. 1b. Class A represents well-formed blastoids with a cavity and adequate ICM embedded in a complete circular TE layer. Class B represents blastoids with a cavity and a small or absent ICM, but a complete TE layer. Class C represents blastoids with an adequate ICM but an irregular or broken TE layer. Class D represents failed blastoids with no cavity, resembling cellular debris, and Class W represents empty microwells where no cells were seeded. Image labeling was performed using the website(https://www.makesense.ai).

### Model Training Methods

Several deep learning models, including ResNet, EfficientNetV2-s and Vision Transformer, were pre-trained on the ImageNet dataset and fine-tuned using our dataset. After evaluating their performance, we selected ResNet18, which demonstrated the best balance between accuracy and throughput. A total of 2107 images were randomly divided into a training set with 1796 data pairs (85%) and a validation set with 311 data pairs (15%), respectively. To diversify the training data and further improve the model performance and generalization, we adopt various data augmentation strategies during the training stage, including horizontal flip, vertical flip, color jitter, and random erasing. The probability for adopting color jitter and random erasing is set to 40% and 20%, respectively. We employ the standard cross entropy loss and the AdamW optimizer to train our model for 36 epochs with a learning rate of 3e-4 and a weight decay of 1e-3. We clip the gradient to stabilize the training and scale the learning rate of the last fully connected layer with 10 to achieve fast convergence. We selected the model trained with 36 epochs to ensure convergent.

### Software

Pipeline Pilot Server 2018 or 2021 (BIOVIA, (https://www.3ds.com/products-services/biovia/products/data-science/pipeline-pilot/)) was used to manage image directories and their associated metadata. All the model was trained on a NVIDIA A100 80GB GPU. We use Anaconda to organize the experiment environment, including Python 3.10.13, PyTorch 2.1.2, CUDA 12.2, and cuDNN 8.9.2.

### Immunofluorescence staining

The samples were first washed with PBS and fixed in 4% paraformaldehyde (PFA) for 15 minutes. Following fixation, the samples were washed three times with PBS, then permeabilized and blocked using a solution containing 0.2% Triton-X-100 and 6% normal donkey serum for 1 hour. The same solution was used to dilute the primary antibodies according to the manufacturer’s recommendations, and the samples were incubated overnight. The primary antibodies used were specific to SOX-2 (Santa Cruz, sc13720, Goat), NANOG (Santa Cruz, sc134218, Mouse), and OCT-4 (CST, 2840p, Rabbit). After primary antibody incubation, the samples were washed three times and then incubated with corresponding secondary antibodies for 1 hour. Nuclear staining was performed using Hoechst 33342 (Beyotime, 1:2,000) for 15 minutes. Filamentous actin (F-actin) was labeled using Acti-stain 488 phalloidin or Acti-stain 555 phalloidin (1:200; Cytoskeleton). This thorough immunofluorescence staining protocol ensures accuracy and reliability, which is essential for subsequent microscopic analysis. Finally, images were captured using a Leica SP8 or Stellaris 8 confocal microscope and processed using Fiji software.

### Preparation of DMSO and LPA gradient dosage

To prepare the medium supplemented with DMSO, first ensure that all materials and reagents are sterile. Dilute the DMSO to the desired final concentration of 0.1% by adding the appropriate volume of DMSO directly to the medium under sterile conditions. Gently mix the medium to ensure even distribution of the DMSO, then filter-sterilize the final solution using a 0.22 µm filter. The prepared medium can be stored at 4°C for short-term use.

For preparing a medium with gradient concentrations of Lysophosphatidic Acid (LPA) at 0 µM, 0.5 µM, 1 µM, 2.5 µM, and 5 µM, begin by preparing a stock solution of LPA at a concentration of 10 mM. Ensure all solutions are sterile and prepare the base medium under sterile conditions. For each concentration, calculate the volume of LPA stock needed based on the desired final concentration. Add the appropriate amount of LPA stock to aliquots of the base medium, mixing thoroughly after each addition to ensure homogeneity. For instance, to prepare 1 mL of 0.5 µM LPA medium, add 0.05 µL of 10 mM LPA stock solution, and follow similar calculations for the other concentrations. Filter-sterilize the solutions if necessary and store the prepared media at 4°C, protected from light, for short-term use.

### Statistical analysis

Statistical analyses in this study were performed with a high degree of rigor and standardization using GraphPad Prism v8.4.2. To determine significant differences between experimental groups, one-way ANOVA was employed, followed by Tukey’s multiple comparisons test for datasets involving three or more groups. For comparisons between two groups, an unpaired, two-tailed Student’s t-test was used. The specific statistical tests applied for each analysis are noted in the corresponding figure legends.

All quantitative data are presented as mean values ± standard error of the mean (SEM), unless otherwise stated. To ensure the reliability and reproducibility of results, each experiment was conducted with at least three independent replicates. Micrographs shown are representative of these replicates. Figures include exact P values to highlight the statistical significance of the findings. Additionally, to enhance clarity and transparency, details regarding sample sizes and the specific statistical tests employed for each experiment are thoroughly described in the figure legends.

## Data and code availability

The datasets used for training and evaluating the models can be accessed and downloaded from ZENODO and is accessible as of the date of publication.

All source code has been deposited in a publicly available repository on GitHub and is accessible as of the date of publication.

## Acknowledgments

This work was financially supported in part by funding from King Abdullah University of Science and Technology (KAUST) - Center of Excellence for Generative AI, under award number 5940 (PW) and by funding from King Abdullah University of Science and Technology (KAUST) – KAUST Center of Excellence for Smart Health (KCSH), under award number 5932 (ML). The work in the group of ML was supported by the KAUST Office of Sponsored Research (OSR) under Award No. BAS/1/1080-01 (ML). We thank J.X. and D.C for administrative support.

## Author contributions

Y.J. performed the experiments related to the blastoid formation and imaging; Z.Fan, Y.J, A.P.C, I.M.S and A.S performed experiments related to establishment of dataset; Yiqing performed experiments related to efficiency of LPA gradient test; Y.Z and M.W assisted Z.Fan in data processing and analysis; Y.T assisted with experimental design and planning; X.Z and Yiqing assisted Z.Fan in figure drawing; Z.Fan, Y.J, A.P.C and I.M.S did blastoid classification and validation; A.S, M.W, X.Z, Y.T and Y.Z provided comprehensive advice in this research; Z.Yu and P.W established and trained the deep learning model; Z.Yu did the optimization and characterization the AI model; Z.Fan, Z.Yu, M.L and P.W wrote the manuscript; Z.Fan and M.L. conceived the study; P.W. and M.L. supervised the study.

