## Supplemental information for "deepBlastoid: A Deep Learning-Based High-Throughput Classifier for Human Blastoids Using Brightfield Images with Confidence Assessment"

### Supplementary Information

| Layer | Height | Width | Depth | Filter Height | Filter Width |
|---|---|---|---|---|---|
| Input | 300 | 300 | 1 | - | - |
| Conv1 | 150 | 150 | 64 | 7 | 7 |
| Max Pool | 75 | 75 | 64 | 3 | 3 |
| ResBlock1 (1st conv) | 75 | 75 | 64 | 3 | 3 |
| ResBlock1 (2nd conv) | 75 | 75 | 64 | 3 | 3 |
| ResBlock2 (1st conv) | 38 | 38 | 128 | 3 | 3 |
| ResBlock2 (2nd conv) | 38 | 38 | 128 | 3 | 3 |
| ResBlock3 (1st conv) | 19 | 19 | 256 | 3 | 3 |
| ResBlock3 (2nd conv) | 19 | 19 | 256 | 3 | 3 |
| ResBlock4 (1st conv) | 10 | 10 | 512 | 3 | 3 |
| ResBlock4 (2nd conv) | 10 | 10 | 512 | 3 | 3 |
| Global Avg Pool | 1 | 1 | 512 | - | - |
| Fully Connected | - | - | 5 | - | - |

**Table1. Key layer information of ResNet-18 convolutional neural network for a 300x300 grayscale image input.**

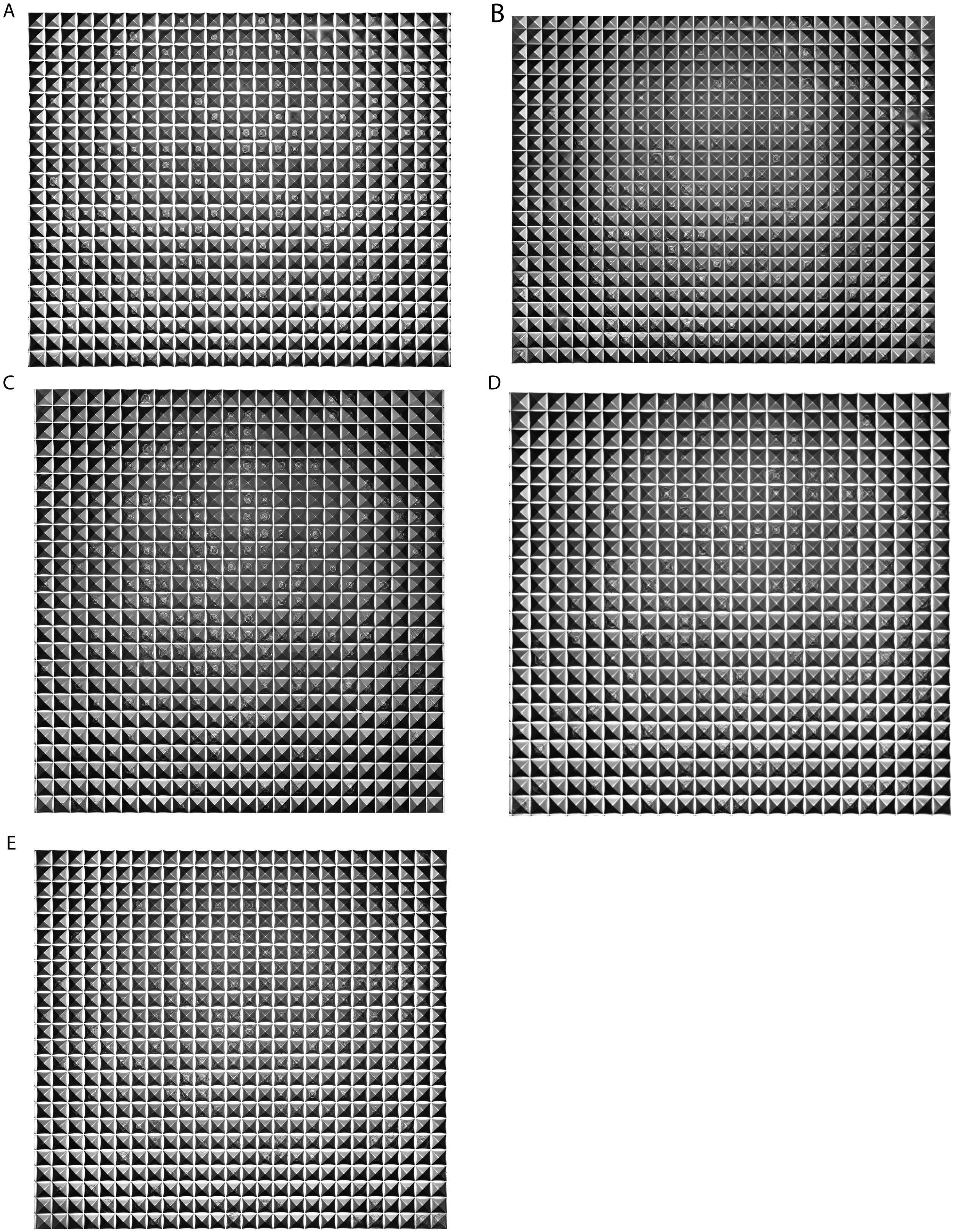
A
B
C
D
E

**Supplementary Fig. 1. Morphology of blastoid in Aggrewell on dose-effect of LPA. (A)**, 0 μM LPA**. (B)**, 0.5 μM LPA**. (C)**, 1 μM LPA**. (D)**, 2.5 μM LPA**. (E)**, 5 μM LPA**.**

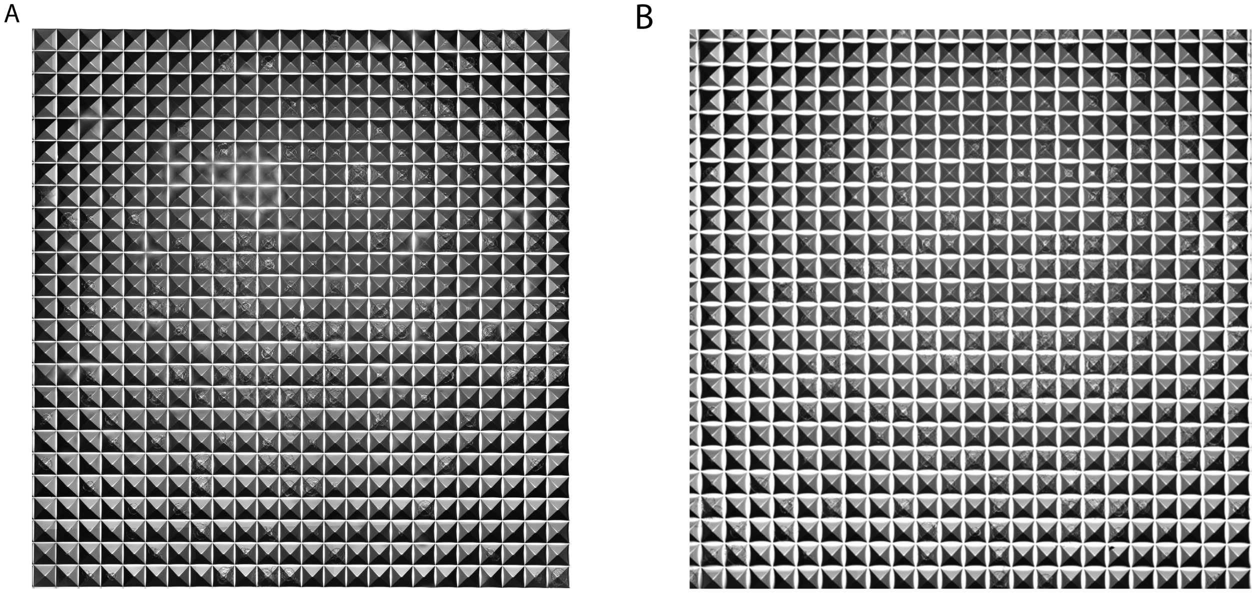

**Supplementary Fig. 2. Morphology of blastoid in Aggrewell on the effect of 0.1%DMSO.**
**(A**), Control group**. (B)**, 0.1%DMSO group**.**
